# Muscle of obese insulin-resistant humans exhibits losses in proteostasis and attenuated proteome dynamics that are improved by exercise training

**DOI:** 10.1101/2023.03.16.532839

**Authors:** Kanchana Srisawat, Connor A Stead, Katie Hesketh, Mark Pogson, Juliette A. Strauss, Matt Cocks, Ivo Siekmann, Stuart M. Phillips, Paulo J. Lisboa, Sam Shepherd, Jatin G Burniston

## Abstract

We examined muscle proteostasis in obese insulin-resistant (OIR) individuals to determine whether endurance exercise could positively influence proteome dynamics in this population. Male OIR (n = 3) and lean, healthy controls (LHC; n = 4) were recruited and underwent a 14-d measurement protocol of daily deuterium oxide (D_2_O) consumption and serial biopsies of vastus lateralis muscle. The OIR group then completed 10-weeks of high-intensity interval training (HIIT), encompassing 3 sessions per week of cycle ergometer exercise with 1 min intervals at 100 % maximum aerobic power (W_max_) interspersed by 1 min recovery periods. The number of intervals per session progressed from 4 to 8, and during weeks 8-10 the 14-d measurement protocol was repeated. The abundance and turnover rates of 880 and 301 proteins, respectively, were measured. OIR and LHC muscle exhibited 352 differences (*p* < 0.05, false discovery rate (*p* < 0.05) differences in protein turnover. OIR muscle was enriched with markers of metabolic stress, protein misfolding and components of the ubiquitin-proteasome system, and the turnover rate of many of these proteins was less compared to LHC muscle. HIIT altered the abundance of 53 proteins and increased the turnover rate of 22 proteins (*p* < 0.05) in OIR muscle and tended to restore proteostasis, evidenced by increasing muscle protein turnover rates and normalizing proteasome composition in OIR participants. In conclusion, obesity and insulin resistance are associated with compromised muscle proteostasis, which can be partially restored by endurance exercise.

## Introduction

The pathogenesis of obesity and type II diabetes is underpinned by defects in metabolic homeostasis, including hyperinsulinaemia and insulin resistance (Kolb *et al*., 2020). Skeletal muscle accounts for up to 80 % of insulin-stimulated glucose uptake in healthy individuals (Baron *et al*., 1988), and a loss in muscle responsiveness to insulin is a central feature of human metabolic disease. In addition to diminishing muscle glucose uptake, insulin resistance contributes to sustained elevations in insulin secretion and chronic hyperinsulinaemia that inhibit adipose tissue lipolysis and hepatic gluconeogenesis. Insulin is also an important regulator of muscle protein turnover (James *et al*., 2017), but there is uncertainty regarding the combined effects of obesity, dyslipidemia and insulin resistance on protein metabolism in human muscle (Freitas & Katsanos, 2022). Chronically elevated insulin levels suppress translocation of glucose transporters to the cell membrane but other aspects of the insulin receptor signalling cascade, including mTORC1 signalling and subsequent effects on protein turnover, may remain stimulated (Kolb *et al*., 2020). Indeed, kinase activity profiling in muscle from healthy lean vs obese insulin-resistant (OIR) individuals (Qi *et al*., 2020) found hyperactivation of JNK stress kinase signalling and hypo-activation of negative regulators of the mTOR pathway in OIR muscle.

The average turnover rate of muscle proteins is lower in obese individuals under fasting conditions (Tran *et al*., 2016; Tran *et al*., 2018) and in the post-absorptive state (Guillet *et al*., 2009). However, when recreationally active lean and obese individuals are studied, there is no difference in the average turnover rate of muscle proteins, either at rest or after a bout of resistance exercise (Hulston *et al*., 2018). When proteins of the myofibrillar fraction are studied, no differences in fractional synthesis rate (FSR) amongst healthy-, over-weight or obese individuals are evident; however, healthy-weight participants exhibit a greater rise in myofibrillar protein FSR after a protein-rich meal (Beals *et al*., 2016). The average FSR of proteins in the muscle mitochondrial fraction is less in the muscle of obese humans compared to normal-weight controls (Guillet *et al*., 2009; Tran *et al*., 2018). However, the protein synthetic response to amino acid provision may be either relatively impaired (Guillet *et al*., 2009) or enhanced (Tran *et al*., 2018) amongst mitochondrial proteins from obese versus normal-weight humans. The various differences in experimental design and the metabolic state of participants in the aforementioned studies make it challenging to reach a consensus on the effects of obesity on muscle protein metabolism. In addition, analyses of mixed-protein data generated from short-duration amino acid tracer studies lack detail on protein-specific responses.

Non-targeted proteomic studies have consistently highlighted decrements to oxidative phosphorylation, a greater reliance on glycolytic metabolism and a shift toward a fast-twitch myofibre profile in the muscle of obese individuals and people with type 2 diabetes (Srisawat *et al*., 2017). Multi-omic investigation, (Vanderboom *et al*., 2022), similarly found transcriptomic and proteomic signatures of impaired mitochondrial function in the muscle of obese individuals, but the down regulation of transcripts relating to protein translation, ribosome and amino acid metabolism in the muscle of obese individuals was not evident at the protein level. The muscle of obese individuals also exhibited blunted and disparate transcriptome, proteome and phosphoproteome responses to acute exercise compared to lean healthy controls. In particular, the transcriptional response to exercise was absent in the muscle of obese participants. Nevertheless, proteins associated with protein translation decreased in abundance and proteins associated with protein degradation increased in abundance, specifically in the muscle of obese participants after exercise (Vanderboom *et al*., 2022). These findings further implicate dysregulation of muscle protein turnover in the context of obesity and indicate that protein-specific responses may occur rather than changes *en masse* to the turnover of all muscle proteins.

While it is challenging to measure the turnover of individual proteins using isotope-labelled amino acid tracers in humans, the stable isotope deuterium oxide (D_2_O) can be readily combined with peptide mass spectrometry to generate synthesis data on a protein-by-protein basis in humans (Burniston, 2019). D_2_O can be administered in the drinking water of free-living humans over days or weeks and is, therefore, less invasive than applications involving isotope-labelled amino acid tracers that require intravenous infusion. The analysis of D_2_O-labelled samples by peptide mass spectrometry and proteomic profiling techniques, generates robust data on the synthesis rate and abundance of each protein (Srisawat *et al*., 2019). Combining protein abundance and turnover data can greatly aid biological interpretation and add a new dimension to muscle analyses (Camera *et al*., 2017). In the current work, we used D_2_O labelling and proteomics to investigate differences in the turnover and abundance of muscle proteins between men that were either lean, healthy individuals or obese individuals with insulin resistance. We hypothesized that obesity is associated with select differences in the turnover, as well as abundance, of proteins in human muscle, and that a programme of aerobic exercise would restore muscle protein homeostasis.

## Methods

### Participants

Men were invited to participate in the study if they were between 30 – 45 years of age and identified as being either overweight/ obese and living a sedentary lifestyle or normal weight and engaged in regular endurance exercise training. Potential participants were given verbal and written details of the study, including potential risks. Inclusion was based on habitual physical activity levels, determined using the Paffenbarger physical activity questionnaire. The potential participants were initially screened by questionnaire and a preliminary health check, including measurement of systolic and diastolic blood pressures, body weight, height, and calculation of body mass index (BMI). Our subsequent inclusion criteria consisted of a reference group of lean, healthy participants (n = 4) and participants with obesity (n = 3). Each participant gave their informed consent to the experimental procedures approved (16/WM/0296) by the Black Country NHS Research Ethics Committee (West Midlands, UK) and conformed with the Declaration of Helsinki, except registration in a clinical trials database.

### Experimental Protocol

Figure 1 provides an overview of the experimental protocol, which consists of a cross-sectional study between OIR and LHC participants at baseline and a longitudinal study of the effect of 10-weeks high intensity interval training (HIIT) in OIR participants only. Anthropological and physiological data, including BMI, body composition, insulin sensitivity and exercise capacity, were collected from all LHC and OIR participants at least 3 d prior to commencing the first 14-day period of D_2_O consumption (i.e. baseline investigation period). Throughout baseline measurements, saliva and blood samples were collected (every day and every 2^nd^ day, respectively), and muscle samples were obtained before D_2_O administration (0 day) and after 4, 9, and 14 days of D_2_O consumption. The LHC group completed the baseline assessment period only, whereas the OIR group undertook a 10-week HIIT intervention. During the final 2 weeks of the HIIT intervention (weeks 8 to 10), the OIR group underwent a second 14-d period of D_2_O consumption, including the collection of saliva, blood and muscle samples. Anthropological and physiological measurements were repeated in OIR participants at least 72 h after completing the 10-week HIIT intervention.

**Figure 1.**
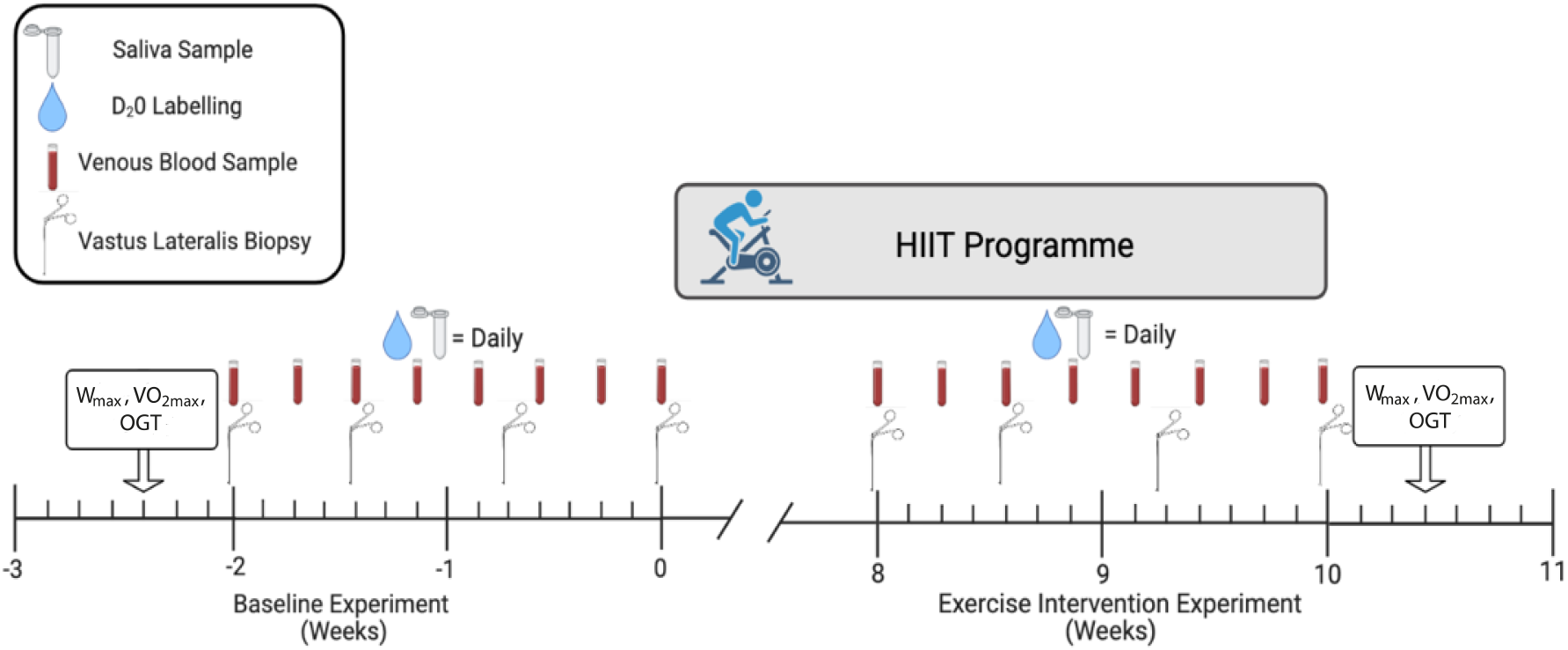
Experimental design and deuterium incorporation. A two-week deuterium oxide (D_2_O) labelling experiment was conducted with lean healthy control (LHC; n = 4) and obese insulin resistant (OIR; n = 3) participants to collect baseline data (weeks −2 – 0). Saliva sampling and D_2_O administration (4 x 50 ml) were conducted daily and venous blood samples were collected every other day. Percutaneous biopsies of vastus lateralis muscle were conducted on days 0, 4, 9 and 14. The OIR group then completed a 10-week high-intensity interval training (HIIT) program. During the last 2 weeks of the training intervention, a repeat of the labelling experiment was conducted to investigate the effects of exercise on the OIR muscle proteome. Physiological data, including peak aerobic power (W_max_), maximum oxygen uptake (VO_2_max) and oral-glucose tolerance (OGT), were measured 48 h prior to the collection of baseline biological samples or 48 h after completion of the 10-week HIIT intervention.

### Assessment of body composition, aerobic exercise capacity, and blood glucose homeostasis

Body composition was measured using whole-body fan-beam dual-energy x-ray absorptiometry (DEXA; Hologic QDR Series, Discovery A, Bedford, MA, USA). Participants were scanned (~180 s) in a supine position, and scans were automatically analysed (QDR software) with manual correction of trunk and limb regions where necessary. Total fat mass (kg), lean mass (kg), and percent body fat (%) are presented as subtotal values excluding head measurements to reduce measurement error.

Following the DEXA scan, participants performed a progressive exercise test to exhaustion on an electronically braked cycle ergometer (Lode BV, Groningen, The Netherlands) to determine their maximal aerobic power (W_max_) and peak oxygen uptake (VO_2peak_). Respiratory gasses were measured using an online gas collection system (Moxus metabolic cart, AEI Technologies, Pittsburgh, Pennsylvania, USA). The test consisted of an initial load of 95 W for 3 min, followed by sequential increments of 35 W every 3 min until cadence was reduced to <50 rpm, at which point the test was terminated. VO_2peak_ was recorded as the highest value obtained during the last 30 s of the test.

On a separate occasion, participants attended the laboratory after an overnight fast (>10 h), having refrained from vigorous exercise in the preceding 48 h period, and underwent an oral glucose tolerance test (OGTT) to determine insulin sensitivity. A resting blood sample (10 ml) was taken before subjects consumed a bolus of 75 g glucose in 250 ml water. Blood samples were collected at 15, 30, 45, 60, 90 and 120 min after glucose consumption. Isotonic saline was used to maintain cannula patency, and blood was collected in serum separator and EDTA-coated vacutainers. Serum and plasma samples were obtained through centrifugation at 1,000 g for 10 min at 4°C and stored at − 80°C for subsequent analysis. Plasma glucose concentrations were determined spectrophotometrically using a glucose oxidase kit and semi-automatic analyser (RX Daytona+; Randox Laboratories, Antrim, UK). Insulin concentrations were determined using a commercially available direct insulin enzyme-linked immunosorbent assay (ELISA) kit (#KAQ1251; Thermo Fisher Scientific, UK). Insulin sensitivity index (ISI) was calculated from fasting plasma glucose and insulin concentrations according to (Matsuda & DeFronzo, 1999; DeFronzo & Matsuda, 2010) and these procedures were repeated after the HIIT in OIR participants.

### HIIT Protocol

The HIIT protocol was similar to that reported by Gillen et al., (Gillen *et al*., 2013). After a 3-min warm-up cycling at 50 W, the OIR group performed repeated cycling bouts at 100 % maximum power output (W_max_) for 60 s, interspersed with 60 s low-intensity recovery cycling at 50 W, maintaining a cadence <50 rpm. Participants trained three times per week for 10 weeks. All participants completed at least 28 (~93 %) of the 30 sessions. Initially, participants performed 4 intervals per training session, which increased by 1 interval after every 2 weeks of training, such that participants performed 8 intervals per training session during weeks 9 and 10.

### Stable isotope labelling *in vivo*

Biosynthetic labelling of newly synthesised proteins was achieved by oral consumption of deuterium oxide (Sigma-Aldrich, UK), consistent with our previous work (Camera *et al*., 2017). Participants consumed 50 ml of 99.8 atom % of D_2_O four times per day (totaling 200 ml per day) approximately 3-4 hours apart, every day over each 14-day labeling period.

### Muscle Biopsy Protocol

Muscle biopsies were taken on day 0, 4, 9 and 14 of each labelling period. All samples were obtained after an overnight fast (> 10 h). Local anesthetic was administered (0.5 % Marcaine) under the skin and over the fascia; samples (~100 mg) of vastus lateralis muscle were taken using the conchotome technique. Muscle samples were blotted to remove excess blood, and visible fat and connective tissue were removed through dissection. Muscle tissue was snap-frozen in liquid nitrogen and stored at −80 °C for subsequent analysis. In total, subjects received two muscle biopsies from each leg in a randomized order over the 14-day experimental periods.

### Calculation of D_2_O Enrichment

During each 14-day period of D_2_O consumption, saliva samples were collected in cryotubes using a passive drool for 60 s prior to the first drink each day. Participants brought saliva samples to the laboratory in cool bags, which were stored at −80 °C for later analysis. A 7 ml venous blood sample was collected from an antecubital vein every 2^nd^ day during D_2_O consumption. Blood samples were collected into serum separator vacutainers, centrifuged at 1,000 g for 10 min at 4°C and stored at −80 °C for subsequent analysis.

Body water enrichment of D_2_O was measured in plasma and saliva samples against external standards constructed by adding D_2_O to PBS over the range from 0.0 to 5.0 % in 0.5 % increments. D_2_O enrichment of aqueous solutions was determined by gas chromatography-mass spectrometry after exchange with acetone (McCabe *et al*., 2006). Samples were centrifuged at 12,000 g, 4°C for 10 min, and 20 μl of plasma supernatant or standard was reacted overnight at room temperature with 2 μl of 10 M NaOH and 4 μl of 5% (v/v) acetone in acetonitrile. Acetone was then extracted into 500 μl chloroform, and water was captured in 0.5 g Na_2_SO_4_ before transferring a 200 μl aliquot of chloroform to an auto-sampler vial. Samples and standards were analysed in triplicate using an Agilent 5973 N mass selective detector coupled to an Agilent 6890 gas chromatography system (Agilent Technologies, Santa Clara, CA, USA). A CD624-GC column (30 m 30.25 mm 31.40 mm) was used in all analyses. Samples (1 μl) were injected using an Agilent 7683 auto sampler. The temperature program began at 50°C, increased by 30°C/min to 150°C and was held for 1 min. The split ratio was 50:1 with a helium flow of 1.5 ml/min. Acetone eluted at ~3 min. The mass spectrometer was operated in the electron impact mode (70 eV), and selective ion monitoring of m/z 58 and 59 was performed using a 10 ms/ ion dwell time.

### Muscle processing

Proteins were extracted from muscle samples as previously described (Camera *et al*., 2017; Hesketh *et al*., 2020). Muscle samples were ground in liquid nitrogen, then homogenized on ice in 10 volumes of 1 % Triton X-100, 50 mM Tris, pH 7.4 (including complete protease inhibitor; Roche Diagnostics, Lewes, United Kingdom) using a PolyTron homogenizer. Homogenates were incubated on ice for 15 min, then centrifuged at 1000 x *g*, 4 °C, for 5 min to fractionate myofibrillar (pellet) from soluble (supernatant) proteins. Soluble proteins were decanted and cleared by further centrifugation at 12,000 x *g*, 4 °C, for 45 min. Myofibrillar proteins were resuspended in a half-volume of homogenization buffer and centrifuged at 1000 x *g*, 4 °C, for 5 min. The washed myofibrillar pellet was then solubilized in lysis buffer (7 M urea, 2 M thiourea, 4% CHAPS, 30 mM Tris, pH 8.5) cleared by centrifugation at 12,000 x *g*, 4 °C, for 45 min. Protein concentrations of the myofibrillar and soluble protein fractions were measured by Bradford assay. Aliquots containing 500 μg protein were precipitated in 5 volumes of ice-cold acetone and incubated for 1 h at −20 °C, and proteins were resuspended in lysis buffer to a final concentration of 5 μg/ μl.

Tryptic digestion was performed using the filter-aided sample preparation (FASP) method (Wisniewski *et al*., 2009). Aliquots containing 100 μg protein were washed with 200 μl of UA buffer (8 M urea, 100 mM Tris, pH 8.5). Proteins were incubated at 37 °C for 15 min in UA buffer containing 100 mM dithiothreitol, followed by incubation (20 min at 4 °C) protected from light in UA buffer containing 50 mM iodoacetamide. UA buffer was exchanged for 50 mM ammonium bicarbonate, and sequencing-grade trypsin (Promega, Madison, WI, USA) was added at an enzyme-to-protein ratio of 1:50. Digestion was allowed to proceed at 37 °C overnight then peptides were collected in 100 μl 50 mM ammonium bicarbonate containing 0.2 % trifluoroacetic acid. Samples containing 4 μg of peptides were de-salted using C_18_ Zip-tips (Millipore) and resuspended in 20 μl of 2.5 % (v/v) ACN, 0.1 % (v/v) FA containing 10 fmol/ μl yeast alcohol dehydrogenase (MassPrep standard; Waters Corp., Milford, MA).

### Liquid Chromatography-mass Spectrometry of the myofibrillar fraction

Label-free liquid chromatography-mass spectrometry of myofibrillar proteins was performed using nanoscale reverse-phase ultra-performance liquid chromatography (NanoAcquity; Waters Corp., Milford, MA) and online electrospray ionization quadrupole-time-of-flight mass spectrometry (Q-TOF Premier; Waters Corp.). Samples (5 μl corresponding to 1 μg tryptic peptides) were loaded by partial-loop injection on to a 180 μm ID x 20 mm long 100 Å, 5 μm BEH C_18_ Symmetry trap column (Waters Corp.) at a flow rate of 5 μl/ min for 3 min in 2.5 % (v/v) ACN, 0.1% (v/v) formic acid. Separation was conducted at 35 °C via a 75 μm ID x 250 mm long 130 Å, 1.7 μm BEH C_18_ analytical reverse-phase column (Waters Corp.). Peptides were eluted using a nonlinear gradient that rose to 37.5 % acetonitrile 0.1% (v/v) formic acid over 90 min at a flow rate of 300 nl/ min. Eluted peptides were sprayed directly into the mass spectrometer via a NanoLock Spray source and Picotip emitter (New Objective, Woburn, MA). Additionally, a LockMass reference (100 fmol/ μl Glu-1-fibrinopeptide B) was delivered to the NanoLock Spray source of the mass spectrometer at a flow rate of 1.5 μl/ min and was sampled at 240 s intervals. For all measurements, the mass spectrometer was operated in positive electrospray ionization mode at a resolution of 10,000 full width at half maximum (FWHM). Before analysis, the time-of-flight analyser was calibrated using fragment ions of [Glu-1]-fibrinopeptide B from m/z 50 to 1990.

Mass spectra for liquid chromatography-mass spectrometry profiling were recorded between 350 and 1600 m/z using mass spectrometry survey scans of 0.45-s duration with an interscan delay of 0.05 s. In addition, equivalent data-dependent tandem mass spectrometry (MS/MS) spectra were collected from each baseline (day 0) sample. MS/MS spectra of collision-induced dissociation fragment ions were recorded over 50–2000 m/z from the 5 most abundant precursor ions of charge 2+ 3+ or 4+ detected in each survey scan. Precursor fragmentation was achieved by collision-induced dissociation at a high (20–40 eV) collision energy throughout 0.25 s per parent ion with an interscan delay of 0.05 s. Acquisition was switched from MS to MS/MS mode when the base peak intensity exceeded a threshold of 30 counts/s and returned to the MS mode when the total ion chromatogram (TIC) in the MS/MS channel exceeded 50,000 counts/s or when 1.0 s (5 scans) were acquired. To avoid repeated selection of peptides for MS/MS, the program used a 30-s dynamic exclusion window.

### Liquid Chromatography-mass Spectrometry of the soluble protein fraction

Data-dependent label-free analysis of soluble protein fractions was performed using an Ultimate 3000 RSLCTM nano system (Thermo Scientific) coupled to a Fusion mass spectrometer (Thermo Scientific). Samples (3 μL corresponding to 600 ng of protein) were loaded on to the trapping column (Thermo Scientific, PepMap100, C_18_, 75 μm X 20 mm), using partial loop injection, for 7 minutes at a flow rate of 9 μL/min with 0.1 % (v/v) TFA. Samples were resolved on a 500 mm analytical column (Easy-Spray C_18_ 75 μm, 2 μm column) using a gradient of 96.2 % A (0.1 % formic acid) 3.8 % B (79.9 % ACN, 20 % water, 0.1 % formic acid) to 50 % A 50 % B over 90 min at a flow rate of 300 nL/min. The data-dependent program used for data acquisition consisted of a 120,000 resolution full-scan MS scan (AGC set to 4^e5^ ions with a maximum fill time of 50 ms) with MS/MS using quadrupole ion selection with a 1.6 m/z window, HCD fragmentation with a normalized collision energy of 32 and LTQ analysis using the rapid scan setting and a maximum fill time of 35 msec. The machine was set to perform as many MS/MS scans as to maintain a cycle time of 0.6 sec. To avoid repeated selection of peptides for MS/MS the program used a 60 s dynamic exclusion window.

### Label-free quantitation of protein abundances

Progenesis Quantitative Informatics for proteomics (Waters Corp.) was used to perform label-free quantitation consistent with our previous work (Camera *et al*., 2017; Hesketh *et al*., 2020; Brown *et al*., 2022). Where appropriate, analytical data were LockMass corrected using the doubly charged monoisotopic ion of the Glu-1-fibrinopeptide B. Prominent ion features were used as vectors to warp each data set to a common reference chromatogram. An analysis window of 15–105 min and 350–1500 m/z was selected. Log-transformed MS data were normalized by inter-sample abundance ratio, and relative protein abundances were calculated using nonconflicting peptides only. Abundance data were then normalised to the 3 most abundant peptides of yeast ADH1 to derive abundance measurements in fmol/ μg protein (Silva *et al*., 2006). MS/MS spectra were exported in Mascot generic format and searched against the Swiss-Prot database (2018.7) restricted to Homo-sapiens (20,272 sequences) using a locally implemented Mascot server (v.2.2.03; www.matrixscience.com). Enzyme specificity was trypsin, which allowed 1 missed cleavage, carbamidomethyl modification of cysteine (fixed). QToF data was searched using m/z errors of 0.3 Da, FUSION data were searched using MS error 10 ppm and MS/MS error 0.6 Da. Mascot output (xml format), restricted to nonhomologous protein identifications, was recombined with MS profile data.

### Measurement of protein synthesis rates

Protein fractional synthesis rates (FSR) were calculated per our previous work (Camera *et al*., 2017). Mass isotopomer abundance data were extracted from MS spectra using Progenesis Quantitative Informatics (Waters Corp.). The abundance of m_0_-m_4_ mass isotopomers was collected over the entire chromatographic peak for nonconflicting peptides used for label-free quantitation. Mass isotopomer information was processed in R version 3.6.2. The incorporation of deuterium into newly synthesized protein causes a decrease in the molar fraction of the peptide monoisotopic (m_0_) peak (Burniston, 2019). Throughout the experiment, changes in mass isotopomer distribution follow a nonlinear bi-exponential pattern due to the rise-to-plateau kinetics in D_2_O enrichment of the body water compartment (measured in plasma samples by GC-MS) and the rise-to-plateau kinetics of D_2_O-labelled amino acids into newly synthesized protein (measured in muscle proteins by LC-MS). Data were fitted using a machine learning approach to optimize for the rate of change in the relative abundance of the monoisotopic (m_0_) peak. The rate of change in mass isotopomer distribution is also a function of the number of exchangeable H sites; this fact was accounted for by referencing each peptide sequence against standard tables that reported the relative enrichment of amino acids by deuterium in humans (Price *et al*., 2012).

### Statistical and bioinformatic Analysis

Statistical analysis was performed in R (Version 3.6.2). Baseline abundance for individual proteins quantified in more than one biopsy were calculated in each participant by taking the median abundance of the protein across the time-series. The post-exercise protein abundances were quantified from the final biopsy only.

Baseline comparisons of participant health/ physiological data (e.g., BMI, MI, VO_2peak_ and, W_max_), protein abundances, and turnover rates between the LHC and OIR groups were analysed using between-subjects ANOVA. Whereas within-subjects ANOVA was used to assess the difference between baseline and post-exercise differences in OIR participants.

Along with a comparison of protein-specific data, the median of the individual protein data of each participant was calculated and used in the statistical analysis to compare the average synthesis rate of all the proteins measured between the groups. Significance was identified as *P* < 0.05. False-discovery rates (q-values; (Storey & Tibshirani, 2003)) were calculated for all protein data to test for false positives. Gene ontology analysis (GO) and protein interactions were investigated using bibliometric mining in the search tool for the retrieval of interacting genes/proteins (STRING) (Szklarczyk *et al*., 2019).

## Results

### Participant health and exercise characteristics

Lean participants engaged in regular aerobic exercise training (~3 sessions of 60 minutes per week) and had a BMI of 24.2 ± 2.4 kg.m^-2^, whereas participants with obesity performed less exercise (< 2 sessions of 30 minutes per week) and had a BMI of 34.0 ± 5.8 kg.m^-2^ (Table 1). Obese participants had a significantly (*P* < 0.01) lower VO_2_peak compared to lean individuals (26.1 ± 4.4 vs 45.5 ± 7.9 ml.kg^-1^.min^-1^, respectively) and significantly lower peak power output (185 ± 26 vs 260 ± 48 W, respectively; *P* = 0.04) during cycle ergometry exercise. The total fat mass of obese participants was 2.4-fold greater than lean participants (*P* = 0.037), which equated to an average 12 % greater (*P* = 0.032) body fat percentage in the participants with obesity. Fasting insulin concentrations tended (*P* = 0.07) to be greater in the obese (32.4 ± 21.6 μIU.ml^-1^) compared to lean participants (8.6 ± 2.9 μIU.ml^-1^) at baseline, but fasting blood glucose concentrations were not different between lean and obese individuals (5.0 ± 0.3 and 5.3 ± 1.4 mmol.l^-1^, respectively; *P* = 0.7). In response to the OGTT, the area under the curve (AUC) for plasma glucose was similar between the obese (893.3 ± 371.0 mmol.l^-1^) and lean (703.3 ± 81.5 mmol.l^-1^; *P* = 0.36) groups, whereas for the AUC of insulin tended (*P* = 0.08) to be greater in obese (13,426 ± 7162 μIU.ml^-1^) compared to lean (5422 ± 2627 μIU.ml^-1^). Participants from the obese group were classified as insulin-resistant based on Matsuda Index (MI; 1.7 ± 0.6 mmol), which was significantly (*P* < 0.01) less than lean participants (5.7 ± 1.4 mmol). Based on the above characteristics, obese insulin resistant (OIR) and lean, healthy controls (LHC) are used throughout the manuscript when referring to data from obese or lean individuals, respectively.

**Table 1.** Participant characteristics and physiological data

|  | LHC<br>(*n* = 4) | OIR (Baseline)<br>(*n* = 3) | OIR (Post-Exercise)<br>(*n* = 3) |
| --- | --- | --- | --- |
| Age (Years) | 38 ± 8 | 38 ± 6 |  |
| Height (cm) | 177.1 ± 5.9 | 179.7 ± 4.1 |  |
| Weight (kg) | 75.6 ± 5.1 | 110 ± 21 | 109 ± 22 |
| BMI (kg.m<sup>2</sup>) | 24.2 ± 2.4* | 34.0 ± 5.8 | 33.8 ± 6.1 |
| Fat mass (kg) | 13.5 ± 3.9* | 32.9 ± 14.11 | 32.4 ± 14.0 |
| Lean mass (kg) | 55.9 ± 3.0** | 70.3 ± 5.7 | 70.6 ± 7.5 |
| Percent body fat (%) | 18.3 ± 4.3* | 30.2 ± 7.5 | 30.1 ± 6.4 |
| VO<sub>2peak</sub> (ml.kg<sup>-1</sup>.min<sup>-1</sup>) | 45.5 ± 7.9** | 26.2 ± 4.4 | 28.6 ± 7.0 |
| Watt<sub>max</sub> (W) | 259.8 ± 48.3* | 185.3 ± 26.0 | 210.3 ± 22.5<sup>#</sup> |
| Fasting glucose (mmol.l<sup>-1</sup>) | 5.0 ± 0.3 | 5.3 ± 1.4 | 5.6 ± 0.6 |
| Glucose AUC (mmol.l<sup>-1</sup>) | 703.3 ± 81.5 | 893.3 ± 371.0 | 783.3 ± 259.6 |
| Fasting insulin (uIU.ml<sup>-1</sup>) | 8.6 ± 2.9<sup>#</sup> | 32.4 ± 21.6 | 29.1 ± 19.5 |
| Insulin AUC (uIU.ml<sup>-1</sup>) | 5422 ± 2627<sup>#</sup> | 13,426 ± 7162 | 12,970 ± 6727 |
| Matsuda Index | 5.7 ± 1.4* | 1.7 ± 0.6 | 2.0 ± 1 |

Physical and physiological characteristics measured during pre- and post-experimental testing of the lean healthy control (LHC) and obese insulin resistant (OIR) individuals at baseline and after 10-weeks HIIT exercise. Data presented as mean ± standard deviation. \*P < 0.05; \*\*P < 0.01; <sup>#</sup>P < 0.1 vs OIR baseline.

After 10-weeks of HIIT, the VO_2_peak of OIR participants increased by 9 % (2.4 ml.kg^-1^.min^-1^; *P* = 0.25), and aerobic peak power increased 14 % (*P* = 0.06) from 185 ± 26 W at baseline to 210 ± 23 W after the HIIT intervention. There was no change in fat mass, lean mass, or body fat percentage (all *P* > 0.05) and exercise training did not significantly alter fasting concentrations of glucose (pre = 5.3 ± 1.4, post = 5.6 ± 0.6 mmol.l^-1^; *P* = 0.57) or insulin (pre = 32.4 ± 21.6, post = 29.1 ± 19.5; *P* = 0.27). The mean glucose and insulin AUC were less (by 110 mmol.l^-1^ and 456 μIU.ml^-1^, respectively, over the 120 min OGTT) in response to OGTT after the 10-week exercise programme, but these improvements did not reach statistical significance (*P* = 0.16 and 0.95, respectively). Similarly, the Matsuda index of OIR participants (2.0 ± 1.0) was 18 % greater than baseline (1.7 ± 0.6) after the HIIT intervention, but this improvement in insulin sensitivity was not statistically significant (*P* = 0.45). Anthropological and physiological data at baseline and after HIIT are presented in Table 1.

### Dynamic proteome profiling of human muscle

Proteomic analysis encompassed 28 muscle samples, including day 0, 4, 9 and 14 time points at baseline (n=4 LHC and n=3 OIR) and during weeks 8-10 of the HIIT intervention (n=3, OIR only). Overall, 1,614 proteins were confidently identified (>1 unique peptide at a false identification threshold of 1 %). After filtering to exclude missing values amongst biological replicates, the abundance of 880 proteins was measured across all sampling times in at least n = 3 participants per group. Protein abundances were stable (Figure 2A) in each participant across each time-series (day 0, 4, 9 and 14) of samples used to investigate the biosynthetic labelling of proteins in each experimental condition. Between groups ANOVA highlighted 352 significant (*P* < 0.05, q < 0.05) differences in protein abundance, including 289 proteins that were more abundant in OIR and 63 that were more abundant in LHC muscle at baseline (Figure 2C). In addition, within-subject ANOVA of day 0 baseline samples and samples were taken after 10 weeks HIIT in OIR participants highlighted 53 statistically significant (*P* < 0.05, q > 0.4) changes in protein abundance, including 33 proteins that increased and 20 proteins that decreased in response to the HIIT intervention (Figure 2F).

**Figure 2.**
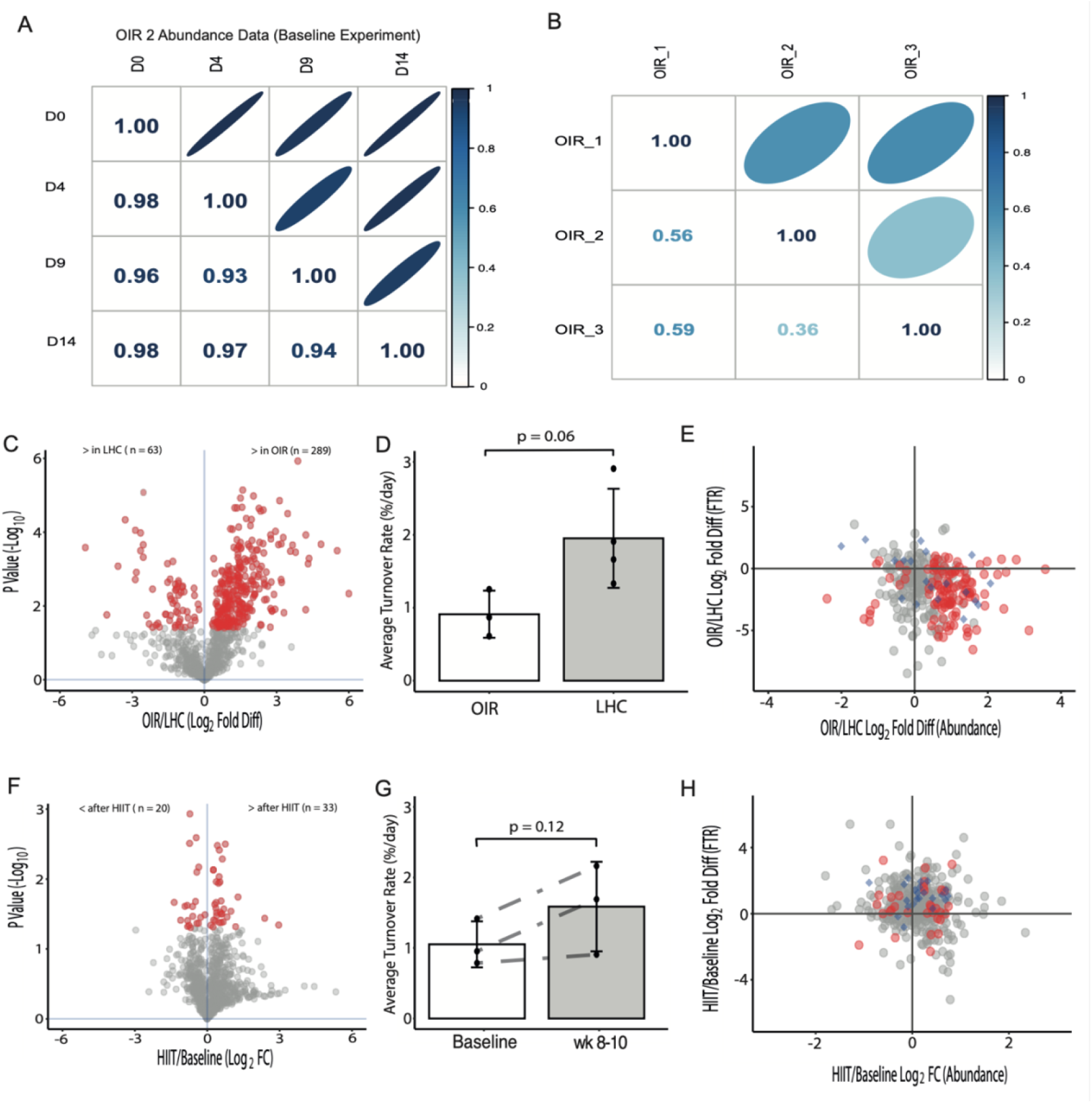
Dynamic proteome profiling of human muscle. **(A)** Representative correlation matrix illustrating the technical reproducibility of muscle protein abundance data (n = 880 proteins quantified) from a participant sampled at days 0, 4, 9 and 14 of the baseline experimental period. **(B)** Representative correlation matrix illustrating the biological variation of protein fractional turnover rates (n = 301 proteins) quantified in n = 3 OIR participants during baseline experimental period. **(C)** Volcano plot illustrating Log_2_ fold-difference in protein abundance between OIR and LHC muscle at baseline (day 0). Statistically significant (p < 0.05) data with a false discovery rate (FDR) < 5 % are highlighted in red. **(D)** Average turnover rate of proteins (n = 301) quantified in OIR (n = 3) and LHC (n = 4) participants during the baseline measurement period. **(E)** Scatter plot of co-occurring differences (Log2 transformed data) in protein abundance (x-axis) and turnover rate (y-axis) in OIR compared to LHC participants. **(F)** Volcano plot illustrating Log_2_ fold-change in protein abundance in OIR participants between baseline (day 0) and the end (day 14) of the HIIT intervention. Statistically significant (p < 0.05) data are highlighted in red, the FDR threshold for this data is >40 %. **(G)** Average turnover rate of proteins (n = 301) quantified in OIR (n = 3) during the baseline measurement period and final two-weeks of the HIIT intervention. **(H)** Scatter plot of co-occurring changes (Log_2_ transformed data) in protein abundance (x-axis) and turnover rate (y-axis) in OIR participants after the IIT intervention.

Deuterium enrichment of body water rose from 0.25 ± 0.06 %/d to a maximum of 3.54 ± 0.5 % during the first 14-day measurement period. By day 0 of the second measurement period, body-water enrichment of deuterium had returned to 0.08 %. During the second 14-day measurement period, deuterium enrichment of body-water rose at a rate of 0.22 ± 0.07 %/d to a maximum of 3.11 ± 0.5 %. There were no significant differences in the rate of body-water enrichment calculated from measurements made using equivalent plasma or saliva samples. High-quality peptide mass isotopomer data were collected for 301 proteins matched across at least *n* = 3 participants in both OIR and LHC groups at baseline or 386 proteins matched across the n=3 OIR participants at baseline and after the HIIT intervention. Protein-specific turnover values were aggregated to derive the average rate of turnover (%/d) of mixed protein, which tended (*P* = 0.061) to be ~2-fold greater in LHC (1.95 ± 0.68) than OIR (0.91 ± 0.32) muscle at baseline (Figure 2D) and increased 1.5-fold (*P* = 0.12) to 1.59 ± 0.63 %/d in OIR muscle during weeks 8-10 of HIIT (Figure 2G).

Nineteen individual proteins exhibited significant (p <0.05, q < 0.04) differences in turnover rate at baseline (Figure 2E), including 11 greater in LHC and 8 greater in OIR. Seven proteins were significantly more abundant in OIR muscle, but their turnover rate was significantly less than in LHC muscle. In addition, 10 proteins exhibited significant differences in the turnover rate but were not different in abundance between OIR and LHC groups (Figure 2H). There were significant (*P* < 0.05) changes in the turnover of 22 individual proteins between baseline and the final 2-weeks of training. Twenty-one proteins changed in turnover independent of changes in protein abundance (19 increasing, 2 decreasing in turnover rate) (Figure 5d). Whereas only 1 protein, 14-3-3 protein Epsilon (14-3-3E), increased in abundance and FSR in response to the HIIT.

Proteins that exhibited significant differences between LHC and OIR baseline or changes in OIR from pre-to post-HIIT were enriched for KEGG pathways relating to energy metabolism (Figure 3), proteasome and cell stress (Figure 4), and the patterns of proteodynamics for each of these biological collections is presented below.

**Figure 3.**
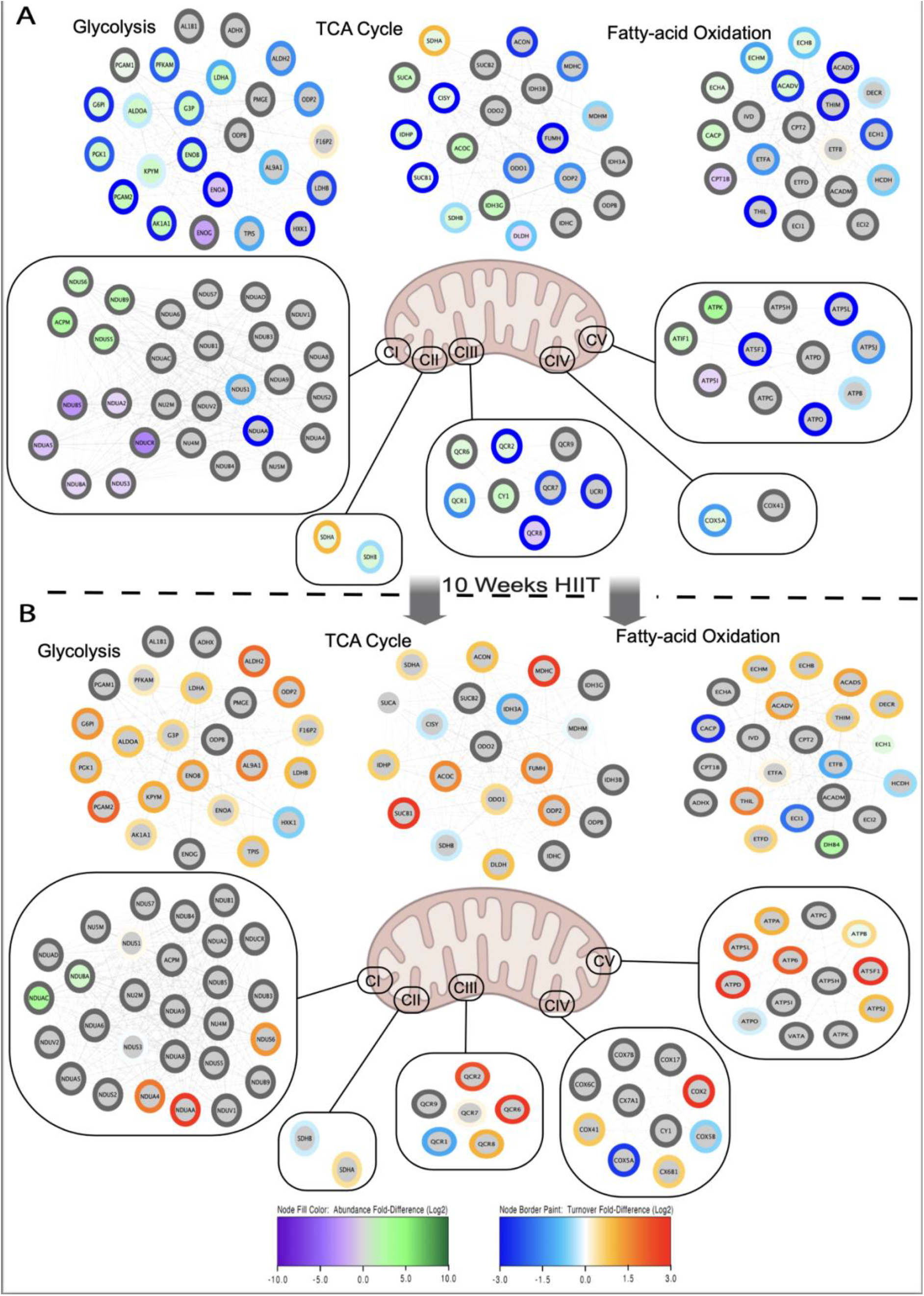
Dynamic proteome profiling of muscle energy metabolism pathways. Nodes represent proteins organized to their principal energy metabolism pathway in muscle and are annotated by their UniProt knowledgebase identifier. **(A)** Node fill colour represents Log2 fold-difference in abundance and node boarder colour represents Log_2_ fold-difference in fractional turnover rate (FSR) between obese-insulin resistant (OIR) and lean healthy control (LHC) participants at baseline. **(B)** Node fill colour represents Log_2_ fold-change in abundance and node boarder colour represents Log_2_ fold-chance in fractional synthesis rate in obese-insulin resistant (OIR) after the 10-week high-intensity interval training (HIIT) intervention. Grey borders indicate missing FSR data. CI – CV represent mitochondrial respiratory chain complexes.

**Figure 4.**
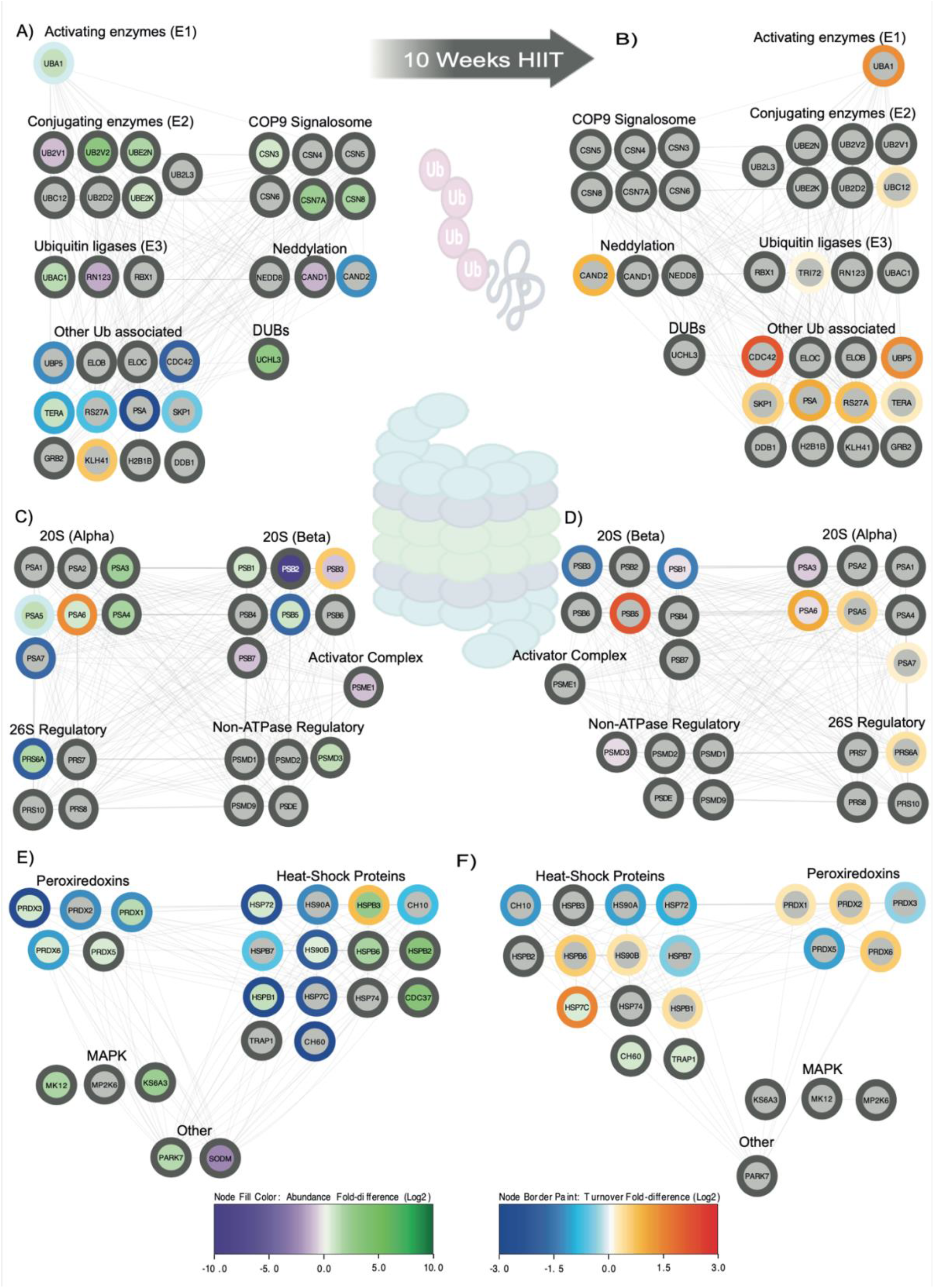
Dynamic proteome profiling of the muscle proteostasis network. Nodes represent proteins annotated by their UniProt knowledgebase identifier and organized to their principal proteostasis network components, including ubiquitin ligase **(A and B)**, proteasome **(C and D)** or heat shock protein and antioxidant system **(E and F)**. **(A, C and E)** Node fill colour represents Log_2_ fold-difference in abundance and node boarder colour represents Log_2_ fold-difference in fractional synthesis rate (FSR) between obese-insulin resistant (OIR) and lean healthy control (LHC) participants at baseline. **(B, D and F)** Node fill colour represents Log_2_ fold-change in abundance and node boarder colour represents Log_2_ fold-chance in fractional turnover rate in obese-insulin resistant (OIR) after the 10-week high-intensity interval training (HIIT) intervention. Grey borders indicate missing FSR data.

### Proteodynamic analysis of proteins associated with muscle energy metabolism pathways

In total, 13 proteins associated with glycolysis/gluconeogenesis (of 24 quantified) were significantly different in abundance between OIR and LHC groups. The majority (11 proteins) of glycolytic proteins were more abundant in OIR muscle. The 2 proteins that were significantly more abundant in LHC were minor muscle isoforms of enolase (α- and γ-enolase), whereas the major muscle isoform (β-enolase; ENOB), was significantly greater in abundance in OIR muscle. Turnover rates were measured for 17 proteins associated with glycolysis/gluconeogenesis and the turnover of each protein tended to be less in OIR than LHC muscle (Figure 3A). In response to HIIT, 2 proteins associated with glycolytic metabolism (glycerol-3-phosphate phosphatase; PGP) and glycogen phosphorylase; PYGM) increased ~1.2-fold in abundance (*P* = 0.038 and 0.030, respectively). Similarly, peroxisomal multifunctional enzyme 2 (DHB4) exhibited a robust increase (~5-fold; *P* = 0.036) in abundance after HIIT (Pre = 4.478 ± 1.408, Post = 23.338 ± 5.469 fmol/μg). The turnover rate of triosephosphate isomerase increased (*P* = 0.046) from 0.26 ± 0.25 %/d at baseline to 0.45 ± 0.31 %/d in trained OIR muscle and the turnover of phosphoglucomutase-1 (PGM1) increased ~3-fold (*P* = 0.011) between baseline (0.34 ± 0.36 %/d) and the final 2-weeks of exercise (0.92 ± 0.46 %/d). In each case, the increases in protein turnover rate were not associated with differences in the abundance of these proteins between baseline and HIIT conditions (Figure 3B).

Significant differences were detected amongst 6 enzymes (of 20 quantified) involved in fatty acid oxidation (FAO) and 9 enzymes (of 21 quantified) of the tricarboxylic acid (TCA) cycle. Most enzymes associated with FAO and TCA cycle were more abundant in OIR muscle (5/6 and 8/9, respectively), whereas dihydrolipoamide dehydrogenase (DLD) was significantly greater in abundance in LHC muscle. Similar to the pattern exhibited by enzymes of glycolytic metabolism, the turnover of the FAO enzymes was generally lower in OIR muscle (Figure 3A). For example, the turnover of enoyl-CoA hydratase (ECHM), was 2.4-fold slower (*P* = 0.044) in OIR (0.60 ± 0.42 %/d) compared to LHC (1.41 ± 0.23 %/d) and ECHM abundance was 1.8-fold greater (*P* = 0.01, q = 0.02) in OIR muscle. The rate limiting enzyme of vitamin B6 metabolism, pyridoxine-5’-phosphate oxidase (PNPO), was 2.5-fold more abundant but had a 16-fold lower turnover rate in OIR (0.24 ± 0.41 %/d) compared to LHC (4.04 ± 2.07 %/d) muscle. The redox enzyme, dehydrogenase/ reductase SDR family member 7 (DHRS7), was 4-fold more abundant (*P* = 0.005, q = 0.01) but had a 2.3-fold (*P* = 0.015) slower turnover rate in OIR compared to LHC muscle at baseline. Exercise led to changes in the abundance or turnover rate of several proteins associated with metabolic pathways (Figure 3B). Long-chain fatty acid CoA ligase-1 (ACSL1) and delta(3,5)-delta(2,4)-dienyol-CoA isomerase, mitochondrial (ECH1) increased in abundance ~1.6 fold (*P* = 0.03 and *P* = 0.27, respectively). Whilst short-chain specific acyl-CoA dehydrogenase (ACADS) did not change in abundance, ACADS increased (*P* = 0.024) 2.4-fold in turnover rate in response to exercise training.

### Proteodynamic analysis of respiratory chain subunits

Overall, 24 proteins (of 61 quantified) belonging to the KEGG pathway “oxidative phosphorylation” (OXPHOS) exhibited significant differences in abundance between OIR and LHC groups (Figure 3A). NADH dehydrogenase (Complex I) exhibited the greatest number of differences and 10 subunits (of 28 quantified) exhibited significant differences in abundance between OIR and LHC groups. Six proteins (NDUB5, NDUS3, NDUCR, NDUA2, NDUA5, and NDUBA) were less abundant in OIR muscle. However, 4 subunits had higher abundance in OIR muscle (NDUB9, NDUS6, NDUS5, and ACPM). Subunits A and B of succinate dehydrogenase (SDH; Complex II) were significantly (*p* < 0.05, FDR < 5 %) more abundant in OIR muscle compared to LHC, but there was no difference in abundance of the 2 membrane-anchoring SDH subunits, C and D. Eight subunits of Complex III (Cytochrome c reductase) were quantified and 5 exhibited significant differences between OIR and LHC participants. Cytochrome c (CY1) and 2 other core subunits of the cytochrome b-c1 complex (QCR1 and QCR2) were significantly more abundant in OIR as was subunit 6 (QCR6) which is associated with the low molecular weight component of Complex III. However, subunit 8 (QCR8), which is also associated with the low molecular weight sub-complex, was significantly less abundant OIR muscle. Cytochrome c oxidase subunit 5A (COX5A) was the only subunit of 7 quantified from Complex IV that was significantly less abundant (*P* = 0.03, q = 0.04) in OIR muscle. Eleven subunits of ATP synthase (Complex V) were analysed, and 3 exhibited significant differences in abundance between LHC and OIR muscle. ATP synthase subunits f (ATPK) and the endogenous inhibitor (ATIF1) were more abundant (8.4-fold and 2.9-fold, respectively), in OIR muscle. Conversely, ATP5I was more abundant in LHC muscle and (1.8-fold). Alongside protein abundance profiling our analysis quantified the turnover rates of 15 OXPHOS subunits. Generally, the turnover data indicated a theme of lower mean turnover rate in OIR muscle (Figure 3) and the rate of turnover of 1 protein, Cytochrome b-c1 complex subunit Rieske (UCRI), was statistically (*P* = 0.025) greater in LHC (2.78 ± 1.20 %/d) compared to OIR (0.37 ± 0.55 %/d) muscle.

Two subunits of respiratory Complex I, including alpha-subcomplex 12 (NDUAC) and beta subcomplex subunit 10 (NDUBA) that were significantly less abundant in OIR compared to LHC at baseline, increased in abundance by 2.4-fold (*P* = 0.018) and 7.8-fold (*P* = 0.045), respectively after HIIT (Figure 3B). ATP synthase subunit beta (ATPB) increased (1.3-fold, *P* = 0.04) from 32.73 ± 9.94 fmol/μg at baseline to 42.27 fmol/μg post-HIIT. Two other ATP synthase subunits (G; ATP5L and A; ATPA) exhibited greater turnover rates after 10 weeks HIIT but their abundance was unaffected. ATP5L exhibited a robust increase (~4-fold, *P* = 0.007) in turnover rate from 0.21 ± 0.34 %/d at baseline to 0.82 ± 0.36 %/d across the final 2-weeks of exercise. ATPA increased >2-fold (*P* = 0.045) in turnover rate in response to training in OIR muscle.

### Proteodynamic analysis of proteins associated with the proteasome, ubiquitination or cellular stress response

In addition to metabolic enzymes, proteins belonging to the KEGG “Proteasome” pathway were highly enriched (q = 5.3e^-4^) amongst the significant differences between OIR and LHC muscle. Several enzymes involved in protein ubiquitination exhibited significantly greater abundance in OIR muscle (Figure 4), including the E1 enzyme, ubiquitin-like modifier-activating enzyme 1 (UBA1), E2 ubiquitin-conjugating enzymes UBE2N, UBE2K, UB2V2 and the E3 ligases, UBAC1 and TRI72. There were some exceptions to this pattern, for example variant 1 of the E2 enzyme, UB2V1, was ~2-fold greater (*P* = 0.003, q = 0.009) in LHC muscle and the abundance of the E3 ligase RNF123, was ~3-fold greater (*P* = 0.002, q = 0.008) in LHC. Three subunits of the COP9 signalosome (a deactivator of Cullin-RING ubiquitin ligases) were more abundant in OIR muscle, and a protein involved in mitophagy, FUN14 domain-containing protein 2, was ~9-fold greater in abundance in OIR (0.3407 ± 0.0920 fmol/μg) than LHC (0.0374 ± 0.0424 fmol/μg).

Our analysis encompassed 24 of the 43 known subunits of the 26S proteasome, including all 7 alpha (PSMA) and 7 beta (PSMB) subunits that make up the 20S catalytic core, 4 subunits of the 19S base region (PSMC), 5 subunits of the 19S lid region (PSMD), and one subunit of the 11S proteasome activator (PSME). Non-ATPase regulatory subunit 3 (PSMD3) and proteasome regulatory subunit 6A (PRS6A) were significantly more abundant in OIR muscle, alongside greater abundances of alpha subunits 3, 4, 5, and 6, and beta subunits 1 and 5, of the core particle (Figure 4C). Conversely beta subunits 2, 3, and 7, and the proteasome activator complex subunit 1 (PSME1) were significantly more abundant in LHC muscle. Protein turnover rates were quantified for 6 proteasome subunits but no statistically significant differences in turnover were identified between OIR and LHC groups.

Fifteen proteins associated with response to stress and chaperone function differed in abundance between OIR and LHC muscle, including 4 isoforms of peroxiredoxin (PRDX), each more abundant in OIR than LHC muscle (Figure 4E). PRDX1 exhibited the greatest difference and was ~2.5-fold greater (*P* = 0.008, q = 0.02) in OIR muscle. Similarly, PRDX3, PRDX5, and PRDX6 were each ~1.5-fold more abundant in OIR muscle. Parkinson disease protein 7 (PARK7) was 2.5-fold greater (*P* < 0.001, q = 0.001) in abundance in OIR muscle (26.89 ± 0.22 fmol/μg) than LHC (10.54 ± 1.80 fmol/μg), whereas the mitochondrial superoxide scavenging enzyme, superoxide dismutase (SODM) was greater (*P* < 0.001, q = 0.003) in abundance in LHC (1.965 ± 0.300 fmol/μg) than OIR muscle (0.3132 ± 0.1259 fmol/μg).

Several proteins associated with maintaining proteostasis were more abundant but had a slower turnover rate in OIR muscle. Heat-shock 70 kDA protein 1 (HSP72) exhibited a > 5-fold slower turnover (*P* = 0.01) in OIR (0.74 ± 0.56 %d) compared to LHC muscle (4.11 ± 1.14 %/d) whilst being ~1.5-fold more abundant (*P* = 0.017, q = 0.028) in OIR. Similarly, the detoxifying enzyme (aldo-keto reductase, mitochondrial; AK1A1) exhibited a >3-fold greater abundance in OIR muscle but turned over at a rate of 1.11 ± 0.31 %/d in OIR and 8.50 ± 1.18 %/d in LHC muscle (7.66-fold slower in OIR; *P* < 0.001). Mitochondrial aldehyde dehydrogenase 2 (ALDH2) is the second major enzyme associated with alcohol metabolism and protects against oxidative stress. ALDH2 also had a significantly greater turnover rate in endurance trained muscle (4.12 ± 1.14 %/d) compared to OIR (1.44 ± 1.18 %/d), however no difference in abundance was identified. Whereas the plasma protein Hemopexin (HEMO), which protects against heme-mediated oxidative stress was 3-fold greater in abundance (*P* = 0.009, q = 0.19) and 2-fold greater in FSR (*P* = 0.011) in OIR than LHC muscle at baseline.

Following the HIIT intervention, 4 proteasome subunits that were more abundant in OIR muscle than LHC under baseline conditions became significantly less abundant after exercise (Figure 4D). Ten weeks of HIIT also led to robust changes in the abundance and/or turnover rates of proteins associated with chaperone functions (Figure 4F). The abundance of the mitochondrial heat shock protein, HSP 75 kDa (TRAP1), increased 1.5-fold (*P* = 0.021) from 0.096 ± 0.030 fmol/μg at baseline to 0.143 ± 0.026 fmol/μg after 10 weeks HIIT. Similarly, chaperonin 60 (CH60) exhibited a 1.6-fold increase (*P* = 0.041) in abundance after the HIIT intervention from 2.790 ± 0.190 fmol/μg at baseline to 4.431 ± 0.665 fmol/μg post-training. The adapter protein 14-3-3E, which may positively regulate the heat shock response increased in abundance from 8.178 ± 1.891 fmol/μg to 9.029 ± 2.218 fmol/μg post-exercise (*P =* 0.046). Notably, the abundance of the chaperone, heat shock cognate 71 kDa (HSP7C) increased 1.4-fold (*P* = 0.042) after HIIT specifically in the myofibrillar fraction. The abundance of HSP7C within the soluble fraction remained stable between pre- and post-exercise in OIR muscle, whereas HSP7C turnover increased 2.7-fold (*P* = 0.004). The chaperone HS90-beta also significantly increased in turnover rate (*P* = 0.046) between baseline (4.46 ± 1.07 %/d) and in response to exercise (5.74 ± 1.23 %/d). In addition, exercise training increased the turnover rates of PRDX2 and ALDH2, which exhibited a significantly greater FSR (*P* = 0.050 and 0.037, respectively) in trained OIR muscle (0.58 ± 0.25 and 5.58 ± 1.87 %/d, respectively) in comparison to baseline (0.43 ± 0.26 and 1.44 ± 1.18 %/d) (Figure 4F).

## Discussion

We have used dynamic proteome profiling to report novel differences in both the abundance and turnover rate of proteins in the muscle of LHC and OIR humans. Our findings point to dysregulation of proteostasis in OIR individuals, while longitudinal analysis of OIR muscle after a 10-week programme of HIIT revealed some restoration of muscle proteostasis. Our data complement and extend knowledge from earlier protein abundance profiling studies, and our application of stable isotope labeling *in vivo* afforded new insight into the dynamic state of proteins in the muscle of OIR participants. Many of the proteins that were more abundant in OIR muscle at baseline exhibited slower turnover rates compared to the muscle of LHC participants. This pattern may indicate a poorer quality of proteins in OIR muscle and point to a loss of muscle proteostasis. Indeed, the fundamental components of the proteostasis network, including the ubiquitin proteasome system (UPS) and heat-shock protein (HSP) chaperones, featured prominently amongst the differences between OIR and LHC muscle proteomes.

We discovered differences in the abundance and turnover rate of UPS components, including the 20S core proteasome, 19S regulatory particle, 11S proteasome activator, ubiquitin (E1, E2 and E3) ligases and components of super-complexes (e.g. Cop9 signalosome) that regulate protein ubiquitination and degradation. E3 ubiquitin ligases underpin the selectivity of UPS-mediated protein degradation and have been a focus of previous mechanistic studies. The E3 ubiquitin ligase, tripartite motif-containing protein 72 (TRIM72), was more abundant and had a lower turnover rate in OIR muscle. TRIM72 ubiquitinates the insulin receptor and insulin receptor substrate-1 (IRS1) and negatively affects muscle insulin signalling (Song *et al*., 2013). Consistent with our findings, muscle TRIM72 abundance is greater in models of obesity and insulin resistance, whereas knock-down of TRIM72 protects against muscle insulin resistance induced by a high-fat diet (Hu & Xiao, 2018). HIIT did not alter TRIM72 abundance but did significantly increase TRIM72 turnover, which may be an early indication of a beneficial effect of exercise training. OIR muscle also had a greater abundance of the E3 ubiquitin ligase RNF123, which is the catalytic subunit of the Kip1 ubiquitin promoting complex (KPC). The KPC is responsible for the degradation of Kip1 (cyclin dependent kinase inhibitor 1B) (Kamura *et al*., 2004) and is an acknowledged regulator of the cell cycle that may also protect against stress-induced apoptosis in striated muscle (Yuan *et al*., 2019).

OIR muscle also exhibited differences in the Cop9 signalosome, which regulates the large family of cullin-RING ubiquitin E3 ligases (CRL) by removing the nedd8 ubiquitin-like modifier from cullin subunits (Mosadeghi *et al*., 2016). Only Nedd8-modified CRL complexes are catalytically active and 3 subunits of the COP9 signalosome (CSN3, CSN7a and CSN8) were more abundant in OIR muscle at baseline (Figure 4), which may indicate lesser activation of CRL enzymes. Cullin-associated NEDD8-dissociated protein 2 (CAND2) is also specific to striated muscle and suppresses the activity of SCF (Skp1-Cullin1-F-box protein)-like ubiquitin E3 ligase complexes by binding culin1 to prevent neddylation (Shiraishi *et al*., 2007). No difference in CAND2 abundance was detected, but the turnover of CAND2 was lesser in OIR compared to LHC at baseline and increased in OIR muscle after exercise training. In addition, the deubiquitinating enzyme, ubiquitin carboxyl-terminal hydrolase isozyme L3 (UCHL3), which hydrolyzes the peptide bond of both ubiquitin and nedd8 modifications (Wada *et al*., 1998), was significantly more abundant in OIR muscle. Together these findings indicate disruption to nedd8 post-translational modifications that regulate the activity of key E3 ligase families in muscle.

Ubiquitin E2 enzymes (UBE2-N, K, V1 and V2) also exhibited different abundances between OIR and LHC muscle, particularly those associated with the regulation of K^63^-polyubiquitination. Ubiquitin-conjugating enzyme E2 N (UBE2N) forms heterodimers with either UBE2 variant 1 (UBE2V1) or UBE2 variant 2 (UBE2V2) and regulates the assembly of K^63^-polyubiquitin chains; whereas UBE2K is responsible for generating branched chains containing both k^48^- and k^63^-linked ubiquitins. Polyubiquitin chains joined at ubiquitin K^48^ are an acknowledged degradative signal, whereas the inclusion of K^63^ linkages may counter the signal for proteasomal degradation (Yang *et al*., 2014). The UBE2N/V2 heterodimer (each more abundant in OIR muscle) is associated with protection against DNA damage (Andersen *et al*., 2005) whereas only UBE2V1 isoform was enriched in LHC muscle. UBE2V1 modulates ubiquitin proteasome responses to proteotoxic stress (Xu *et al*., 2020) and the greater likelihood of UBE2N/UBE2V1 heterodimers in LHC muscle may be associated with greater proteome stability. Conversely, UBE2V2 can be modified by reactive electrophiles and may lead to hyperactivation of UBE2N to promote K^63^-polyubiquitination and genome protection (Zhao *et al*., 2018). However, UBE2K was also significantly more abundant in OIR muscle and is responsible for the formation of branched polyubiquitin chains that contain K^48^-as well as K^63^-linkages (Pluska *et al*., 2021). UBE2K may enhance proteasomal degradation of proteins carrying K^63^-polyubiquitin chains (Ohtake *et al*., 2018). These findings highlight a complex interplay between E2 ligases and suggest the distribution of K^48^- and K^63^-linked polyubiquitin chains was altered in the muscle of OIR participants.

The catalytically active subunits of the core proteasome (beta 1, 2 and 5) differed in abundance between OIR and LHC muscle. The beta-1 and beta-5 subunits were significantly more abundant in OIR muscle, whereas the beta-2 subunit was significantly less abundant than LHC muscle; four alpha ring subunits were also more abundant in OIR muscle. Previous studies (Hwang *et al*., 2010; Vanderboom *et al*., 2022) similarly report some but not all proteasome subunits exhibit differences amongst the muscle of lean, obese and T2DM patients. Currently, it is uncertain whether the abundance of individual proteasome subunits measured in muscle homogenates reflects the activity of the proteasome (Jenkins *et al*., 2020). Proteasome activity is also modified by changes to regulatory subunits, including the 11S proteasome activator (PA28α; PSME1), which increases specifically in the muscle of obese participants after acute exercise (Vanderboom *et al*., 2022). Similarly, we found PSME1 was more abundant in OIR muscle after 10-weeks HIIT (Figure 4D) and our earlier analysis of rat heart responses to exercise (Burniston, 2009) also demonstrated that endurance training increases the abundance of the PSME1 subunit. Overexpression of PA28α is associated with increased degradation of oxidatively damaged proteins in rat neonatal ventricular myocytes (Li *et al*., 2011). Whereas streptozotocin-induced insulin-dependent diabetes is associated with reduced muscle PA28α content and loss of proteasome activity (Merforth *et al*., 2003). Furthermore, PA28-null mice exhibit hepatic steatosis, decreased hepatic insulin signaling, and increased hepatic glucose production (Otoda *et al*., 2013). Therefore, despite ambiguous differences in catalytically active subunits of the core proteasome, 10-weeks HIIT likely improved the capacity for proteasomal degradation in OIR muscle via 11S proteasome activation.

Heat shock proteins (HSP) are the second major constituents of the proteostasis network and are widely-acknowledged components of muscle responses to exercise. HSP are categorized based on their molecular weight into major families and the 90 kDa-, 70 kDa- and small (< 45 kDa) heat-shock proteins. Small heat shock proteins (sHSP) exhibited differences in abundance between OIR and LHC muscle that were consistent with previous literature. For example, HSP27 (HSPB1) was significantly more abundant in OIR and is also more abundant in the muscle of Goto-Kakizaki rats (Mullen *et al*., 2011) and in myoblasts generated from the muscle of type 2 diabetic patients (Al-khalili *et al*., 2013). HSPB1 and HSPB6 (HSP20) are well studied in the context of skeletal muscle responses to exercise and each of these proteins exhibited higher rates of turnover after 10-weeks HIIT (Figure 4F). Small HSP (sHSP) function in homo- or hetero-oligomers of various sizes and complexity and the observed differences across several sHSP (Figure 4E) may indicate changes to the size or composition of sHSP oligomers.

sHSP bind efficiently with misfolded proteins but lack ATPase activity and cannot (re-) fold substrate proteins directly (Haslbeck *et al*., 2019). Therefore, sHSP work cooperatively with other chaperone complexes, e.g. by preparing proteins for refolding by HSP70 (Goncalves *et al*., 2021). The inducible HSP72 and the constitutively expressed heat shock cognate (HSP7C) were each more abundant in OIR than LHC muscle. HSP72 abundance increases in human muscle after exhaustive exercise but returns to basal levels within 3 h after the cessation of exercise (Febbraio *et al*., 2002). We report chronic elevation of HSP72 in OIR muscle, which may be evidence of sustained stress and an elevated requirement for refolding damaged proteins (Gupta *et al*., 2010). Indeed, muscle-specific overexpression of HSP72 can protect against the development of insulin resistance induced by a high-fat diet in mice (Chung *et al*., 2008). In the current work, HIIT did not effect HSP72 but did significantly increase the turnover rate of HSP7C, which may support a general improvement in proteome quality by enhancing the capability of HSP7C to orchestrate chaperone-mediated degradative process (Fernández-Fernández & Valpuesta, 2018).

HSP70 complexes may, in turn, pass client proteins to HSP90 complexes, including HSP90-alpha (HS90A) and HSP90-beta (HS90B), which are abundant cytosolic proteins that (re-) fold newly synthesized or incorrectly folded protein clients. Pharmacological inhibition (Lee *et al*., 2013) or knockdown (Jing *et al*., 2018) of HSP90 improves insulin sensitivity in rodent models of diabetes or diet-induced obesity. Consistent with findings in patients with type 2 diabetes (Venojärvi *et al*., 2014), HSP90B was more abundant in OIR muscle (Figure 4E). HIIT did not alter the abundance of either HSP90 isoform but the turnover rate HS90B increased significantly in OIR muscle after 10-weeks of training (Figure 4F). HSP90 function is regulated by post-translational modifications, including oxidation (Backe *et al*., 2020), and a greater turnover of HSP90 in trained muscle may equate to a greater proportion of non-modified HSP90 proteins, which have preserved functional capacity (Beck *et al*., 2012). In addition, the mitochondrially targeted homolog of HSP90, TRAP1, was more abundant in OIR muscle after HIIT, and TRAP1 may offer greater protection against mitochondrial apoptosis induced by reactive oxygen species (Montesano Gesualdi *et al*., 2007).

Redox signalling contributes to the muscle response to exercise, and oxidative stress is a proposed mechanism of muscle dysfunction associated with obesity. PARK7 (DJ-1) and peroxiredoxins (PRDX) were generally more abundant in OIR than LHC muscle. PARK7 is a redox-sensitive chaperone that may reverse methylglyoxal and glyoxal-glycated protein modifications (Richarme *et al*., 2015) that can be elevated in the muscle of obese individuals with type 2 diabetes (Mey *et al*., 2018). PRDX enzymes are the primary scavengers of cellular H2O2 and may underpin the hormesis response of muscle to exercise-induced oxidative stress (Xia *et al*., 2023). Our findings (Figure 4E and F) add to reports that PRDX2 and PRDX6 are more abundant in the skeletal muscle of type 2 diabetic patients (Brinkmann *et al*., 2012), and that PRDX5 in more abundant in myoblasts derived from muscle biopsies of type 2 diabetic patients (Al-khalili *et al*., 2013). We found no change in PRDX abundances after the 10-week HIIT intervention, whereas (Brinkmann *et al*., 2012) reported muscle PRDX5 abundance increases in type 2 diabetic patients after exercise training. PRDX enzymes undergo reversible redox modifications, for example, PRDX3 becomes more oxidized in human muscle during HIIT (Pugh *et al*., 2021). We report that the turnover rate of PRDX3 was relatively low in OIR compared to LHC muscle and further declined after the 10-week HIIT intervention, which may be associated with changes to the modification state and dimerization of PRDX3 in exercised muscle.

Tran et al., (2019) reports the average turnover of mixed protein is less in the muscle of obese compared to lean humans and used targeted analysis of ATPB in ^2^H_10_-leucince labelled samples to report a lesser protein-specific turnover rate of muscle ATB in obese individuals. We also found the protein-specific turnover rate of ATPB was less in OIR muscle, and our non-targeted analysis, which encompassed a further 10 subunits of Complex V, highlighted that subunits AT5F1, ATP5L and ATPO also exhibited lesser rates of turnover in OIR muscle (Figure 3A). When our protein-specific turnover data are aggregated, the average turnover rate of mixed protein tended to be less in OIR than LHC individuals (Figure 2D). However, this pattern was not uniform at the protein-specific level and the turnover of proteins, including kelch-like protein 41 (KLH41), HSPB3 and 2 subunits (PSA6 and PSB3) of the core proteasome tended to be greater in OIR than LHC muscle (Figure 4E). Furthermore, we report a trend towards a greater average protein turnover in OIR muscle undergoing HIIT (Figure 2G) but this response pattern was, again, not uniform at the protein-specific level. In particular, some subunits of the core proteasome, heat shock proteins and peroxiredoxins exhibited lesser turnover rates after HIIT in OIR muscle (Figure 4D and F). Therefore, dynamic proteome profiling adds protein-specific detail to trends observed in mixed protein data and highlights proteins that exhibit responses that are inverse to the overall trend in average turnover of the protein mixture.

Our analysis of protein abundance and turnover responses in human muscle is unique and has yielded new insight into losses in muscle proteostasis associated with obesity. However, we acknowledge our sample size of n = 4 LHC and n = 3 OIR participants limits the extrapolation of our findings to larger populations, and more extensive studies are required to pursue this line of enquiry. Where data exist in the previous literature, our current findings align well with existing knowledge regarding the effects of obesity on skeletal muscle. In agreement with our meta-analysis of protein abundance profiling literature (Srisawat *et al*., 2017), glycolytic enzymes were enriched in OIR muscle and mitochondrial Complex I emerged as a point of convergence between the effects of metabolic disease and exercise training. However, differences in protein-specific turnover rates were the more prominent feature observed across metabolic enzymes between OIR and LHC at baseline (Figure 3A) or in OIR muscle before and after the 10-week HIIT intervention (Figure 3B), which add new information on the effects of obesity in human muscle.

In conclusion, the muscle proteome of obese insulin-resistant humans exhibits widespread evidence of losses in proteostasis and elevated proteome stress characterised by differences in the abundance and turnover rate of heat shock proteins and perturbations to the ubiquitin proteasome system. Ten-weeks of HIIT tended to improve the quality of the proteome by altering proteasome composition and enhancing the turnover rates of metabolic enzymes. We observed changes in the turnover rate of energy metabolism enzymes without exercise-induced changes in protein abundance; therefore, proteodynamic analysis offers new insight into muscle exercise responses. Losses in proteostasis are well-established in age-related diseases, and our discoveries highlight a need to further investigate whether losses in proteostasis also underpin earlier pre-clinical stages of human diseases.

## References

Al-khalili L, Castro TD, Östling J & Massart J. (2013). Profiling of human myotubes reveals an intrinsic proteomic signature associated with type 2 diabetes. Biochemical Pharmacology 2, 25–38.

Andersen PL, Zhou H, Pastushok L, Moraes T, McKenna S, Ziola B, Ellison MJ, Dixit VM & Xiao W. (2005). Distinct regulation of Ubc13 functions by the two ubiquitin-conjugating enzyme variants Mms2 and Uev1A. J Cell Biol 170, 745–755.

Backe SJ, Sager RA, Woodford MR, Makedon AM & Mollapour M. (2020). Post-translational modifications of Hsp90 and translating the chaperone code. J Biol Chem 295, 11099–11117.

Beals JW, Sukiennik RA, Nallabelli J, Emmons RS, van Vliet S, Young JR, Ulanov AV, Li Z, Paluska SA, De Lisio M & Burd NA. (2016). Anabolic sensitivity of postprandial muscle protein synthesis to the ingestion of a protein-dense food is reduced in overweight and obese young adults. Am J Clin Nutr 104, 1014–1022.

Beck R, Dejeans N, Glorieux C, Creton M, Delaive E, Dieu M, Raes M, Leveque P, Gallez B, Depuydt M, Collet JF, Calderon PB & Verrax J. (2012). Hsp90 is cleaved by reactive oxygen species at a highly conserved N-terminal amino acid motif. PLoS One 7, e40795.

Brinkmann C, Chung N, Schmidt U, Kreutz T, Lenzen E, Schiffer T, Geisler S, Graf C, Montiel-Garcia G, Renner R, Bloch W & Brixius K. (2012). Training alters the skeletal muscle antioxidative capacity in non-insulin-dependent type 2 diabetic men. Scand J Med Sci Sports 22, 462–470.

Brown AD, Stewart CE & Burniston JG. (2022). Degradation of ribosomal and chaperone proteins is attenuated during the differentiation of replicatively aged C2C12 myoblasts. J Cachexia Sarcopenia Muscle 13, 2562–2575.

Burniston JG. (2009). Adaptation of the rat cardiac proteome in response to intensity-controlled endurance exercise. Proteomics 9, 106–115.

Burniston JG. (2019). Investigating Muscle Protein Turnover on a Protein-by-Protein Basis Using Dynamic Proteome Profiling. 1 edn, ed. Burniston JG & Chen Y-w, pp. 171–190. Springer, New York.

Camera DM, Burniston JG, Pogson MA, Smiles WJ & Hawley JA. (2017). Dynamic proteome profiling of individual proteins in human skeletal muscle after a high-fat diet and resistance exercise. FASEB J 31, 5478–5494.

Chung J, Nguyen AK, Henstridge DC, Holmes AG, Chan MH, Mesa JL, Lancaster GI, Southgate RJ, Bruce CR, Duffy SJ, Horvath I, Mestril R, Watt MJ, Hooper PL, Kingwell BA, Vigh L, Hevener A & Febbraio MA. (2008). HSP72 protects against obesity-induced insulin resistance. Proc Natl Acad Sci U S A 105, 1739–1744.

DeFronzo RA & Matsuda M. (2010). Reduced time points to calculate the composite index. Diabetes Care.

Febbraio MA, Steensberg A, Walsh R, Koukoulas I, van Hall G, Saltin B & Pedersen BK. (2002). Reduced glycogen availability is associated with an elevation in HSP72 in contracting human skeletal muscle. J Physiol 538, 911–917.

Fernández-Fernández MR & Valpuesta JM. (2018). Hsp70 chaperone: a master player in protein homeostasis. F1000Res 7.

Freitas EDS & Katsanos CS. (2022). (Dys)regulation of Protein Metabolism in Skeletal Muscle of Humans With Obesity. Frontiers in Physiology 13.

Gillen JB, Percival ME, Ludzki A, Tarnopolsky MA & Gibala MJ. (2013). Interval training in the fed or fasted state improves body composition and muscle oxidative capacity in overweight women. Obesity 21, 2249–2255.

Goncalves CC, Sharon I, Schmeing TM, Ramos CHI & Young JC. (2021). The chaperone HSPB1 prepares protein aggregates for resolubilization by HSP70. Sci Rep 11, 17139.

Guillet C, Delcourt I, Rance M, Giraudet C, Walrand S, Bedu M, Duche P & Boirie Y. (2009). Changes in basal and insulin and amino acid response of whole body and skeletal muscle proteins in obese men. J Clin Endocrinol Metab 94, 3044–3050.

Gupta S, Deepti A, Deegan S, Lisbona F, Hetz C & Samali A. (2010). HSP72 protects cells from ER stress-induced apoptosis via enhancement of IRE1alpha-XBP1 signaling through a physical interaction. PLoS Biol 8, e1000410.

Haslbeck M, Weinkauf S & Buchner J. (2019). Small heat shock proteins: Simplicity meets complexity. J Biol Chem 294, 2121–2132.

Hesketh SJ, Sutherland H, Lisboa PJ, Jarvis JC & Burniston JG. (2020). Adaptation of rat fast-twitch muscle to endurance activity is underpinned by changes to protein degradation as well as protein synthesis. FASEB journal: official publication of the Federation of American Societies for Experimental Biology 34, 10398–10417.

Hu X & Xiao RP. (2018). MG53 and disordered metabolism in striated muscle. Biochim Biophys Acta Mol Basis Dis 1864, 1984–1990.

Hulston CJ, Woods RM, Dewhurst-Trigg R, Parry SA, Gagnon S, Baker L, James LJ, Markey O, Martin NRW, Ferguson RA & van Hall G. (2018). Resistance exercise stimulates mixed muscle protein synthesis in lean and obese young adults. Physiol Rep 6, e13799.

Hwang H, Bowen BP, Lefort N, Flynn CR, De Filippis EA, Roberts C, Smoke CC, Meyer C, Højlund K, Yi Z & Mandarino LJ. (2010). Proteomics analysis of human skeletal muscle reveals novel abnormalities in obesity and type 2 diabetes, pp. 33–42.

James HA, O’Neill BT & Nair KS. (2017). Insulin Regulation of Proteostasis and Clinical Implications, pp. 310–323. Elsevier Inc.

Jenkins EC, Shah N, Gomez M, Casalena G, Zhao D, Kenny TC, Guariglia SR, Manfredi G & Germain D. (2020). Proteasome mapping reveals sexual dimorphism in tissue-specific sensitivity to protein aggregations. EMBO Rep 21, e48978.

Jing E, Sundararajan P, Majumdar ID, Hazarika S, Fowler S, Szeto A, Gesta S, Mendez AJ, Vishnudas VK, Sarangarajan R & Narain NR. (2018). Hsp90β knockdown in DIO mice reverses insulin resistance and improves glucose tolerance. Nutr Metab (Lond) 15, 11.

Kamura T, Hara T, Matsumoto M, Ishida N, Okumura F, Hatakeyama S, Yoshida M, Nakayama K & Nakayama KI. (2004). Cytoplasmic ubiquitin ligase KPC regulates proteolysis of p27(Kip1) at G1 phase. Nat Cell Biol 6, 1229–1235.

Kolb H, Kempf K, Röhling M & Martin S. (2020). Insulin: too much of a good thing is bad. BMC medicine 18, 224–224.

Lee J-H, Gao J, Kosinski PA, Elliman SJ, Hughes TE, Gromada J & Kemp DM. (2013). Heat shock protein 90 (HSP90) inhibitors activate the heat shock factor 1 (HSF1) stress response pathway and improve glucose regulation in diabetic mice. Biochemical and Biophysical Research Communications 430, 1109–1113.

Li J, Powell SR & Wang X. (2011). Enhancement of proteasome function by PA28α overexpression protects against oxidative stress. FASEB J 25, 883–893.

Matsuda M & DeFronzo RA. (1999). Insulin sensitivity indices obtained from oral glucose tolerance testing: Comparison with the euglycemic insulin clamp. Diabetes Care 22, 1462–1470.

McCabe BJ, Bederman IR, Croniger C, Millward C, Norment C & Previs SF. (2006). Reproducibility of gas chromatography-mass spectrometry measurements of2H labeling of water: Application for measuring body composition in mice. Analytical Biochemistry 350, 171–176.

Merforth S, Kuehn L, Osmers A & Dahlmann B. (2003). Alteration of 20S proteasome-subtypes and proteasome activator PA28 in skeletal muscle of rat after induction of diabetes mellitus. Int J Biochem Cell Biol 35, 740–748.

Mey JT, Blackburn BK, Miranda ER, Chaves AB, Briller J, Bonini MG & Haus JM. (2018). Dicarbonyl stress and glyoxalase enzyme system regulation in human skeletal muscle. Am J Physiol Regul Integr Comp Physiol 314, R181–R190.

Montesano Gesualdi N, Chirico G, Pirozzi G, Costantino E, Landriscina M & Esposito F. (2007). Tumor necrosis factor-associated protein 1 (TRAP-1) protects cells from oxidative stress and apoptosis. Stress 10, 342–350.

Mosadeghi R, Reichermeier KM, Winkler M, Schreiber A, Reitsma JM, Zhang Y, Stengel F, Cao J, Kim M, Sweredoski MJ, Hess S, Leitner A, Aebersold R, Peter M, Deshaies RJ & Enchev RI. (2016). Structural and kinetic analysis of the COP9-Signalosome activation and the cullin-RING ubiquitin ligase deneddylation cycle. Elife 5.

Mullen E, O’Reilly E & Ohlendieck K. (2011). Skeletal muscle tissue from the Goto-Kakizaki rat model of type-2 diabetes exhibits increased levels of the small heat shock protein Hsp27. Mol Med Rep 4, 229–236.

Ohtake F, Tsuchiya H, Saeki Y & Tanaka K. (2018). K63 ubiquitylation triggers proteasomal degradation by seeding branched ubiquitin chains. Proceedings of the National Academy of Sciences 115, E1401–E1408.

Otoda T, Takamura T, Misu H, Ota T, Murata S, Hayashi H, Takayama H, Kikuchi A, Kanamori T, Shima KR, Lan F, Takeda T, Kurita S, Ishikura K, Kita Y, Iwayama K, Kato K, Uno M, Takeshita Y, Yamamoto M, Tokuyama K, Iseki S, Tanaka K & Kaneko S. (2013). Proteasome dysfunction mediates obesity-induced endoplasmic reticulum stress and insulin resistance in the liver. Diabetes 62, 811–824.

Pluska L, Jarosch E, Zauber H, Kniss A, Waltho A, Bagola K, von Delbrück M, Löhr F, Schulman BA, Selbach M, Dötsch V & Sommer T. (2021). The UBA domain of conjugating enzyme Ubc1/Ube2K facilitates assembly of K48/K63-branched ubiquitin chains. The EMBO Journal 40, e106094.

Price JC, Holmes WE, Li KW, Floreani NA, Neese RA, Turner SM & Hellerstein MK. (2012). Measurement of human plasma proteome dynamics with2H2O and liquid chromatography tandem mass spectrometry. Analytical Biochemistry 420, 73–83.

Pugh JN, Stretton C, McDonagh B, Brownridge P, McArdle A, Jackson MJ & Close GL. (2021). Exercise stress leads to an acute loss of mitochondrial proteins and disruption of redox control in skeletal muscle of older subjects: An underlying decrease in resilience with aging? Free radical biology & medicine, 135907–135907.

Qi Y, Zhang X, Seyoum B, Msallaty Z, Mallisho A, Caruso M, Damacharla D, Ma D, Al-Janabi W, Tagett R, Alharbi M, Calme G, Mestareehi A, Draghici S, Abou-Samra A, Kowluru A & Yi Z. (2020). Kinome Profiling Reveals Abnormal Activity of Kinases in Skeletal Muscle From Adults With Obesity and Insulin Resistance. J Clin Endocrinol Metab 105.

Richarme G, Mihoub M, Dairou J, Chi Bui L, Leger T & Lamouri A. (2015). Parkinsonism-associated protein DJ-1/park7 is a major protein deglycase that repairs methylglyoxal-and glyoxal-glycated cysteine, arginine, and lysine residues. Journal of Biological Chemistry 290, 1885–1897.

Shiraishi S, Zhou C, Aoki T, Sato N, Chiba T, Tanaka K, Yoshida S, Nabeshima Y, Nabeshima YI & Tamura TA. (2007). TBP-interacting Protein 120B (TIP120B)/Cullin-associated and Neddylation-dissociated 2 (CAND2) inhibits SCF-dependent ubiquitination of myogenin and accelerates myogenic differentiation. Journal of Biological Chemistry 282, 9017–9028.

Silva JC, Gorenstein MV, Li G-z, Vissers JPC & Geromanos SJ. (2006). Absolute Quantification of Proteins by LCMS E. Molecular & Cellular Proteomics 5, 144–156.

Song R, Peng W, Zhang Y, Lv F, Wu HK, Guo J, Cao Y, Pi Y, Zhang X, Jin L, Zhang M, Jiang P, Liu F, Meng S, Zhang X, Jiang P, Cao CM & Xiao RP. (2013). Central role of E3 ubiquitin ligase MG53 in insulin resistance and metabolic disorders. Nature 494, 375–379.

Srisawat K, Hesketh K, Cocks M, Strauss J, Edwards BJ, Lisboa PJ, Shepherd S & Burniston JG. (2019). Reliability of Protein Abundance and Synthesis Measurements in Human Skeletal Muscle. PROTEOMICS 1900194, 1900194–1900194.

Srisawat K, Shepherd SO, Lisboa PJ & Burniston JG. (2017). A Systematic Review and Meta-Analysis of Proteomics Literature on the Response of Human Skeletal Muscle to Obesity/Type 2 Diabetes Mellitus (T2DM) Versus Exercise Training. Proteomes 5, 30–30.

Storey JD & Tibshirani R. (2003). Statistical significance for genomewide studies. Proceedings of the National Academy of Sciences of the United States of America 100, 9440–9445.

Szklarczyk D, Gable AL, Lyon D, Junge A, Wyder S, Huerta-Cepas J, Simonovic M, Doncheva NT, Morris JH, Bork P, Jensen LJ & Von Mering C. (2019). STRING v11: Protein-protein association networks with increased coverage, supporting functional discovery in genome-wide experimental datasets. Nucleic Acids Research 47, D607–D613.

Tran L, Hanavan PD, Campbell LE, De Filippis E, Lake DF, Coletta DK, Roust LR, Mandarino LJ, Carroll CC & Katsanos CS. (2016). Prolonged Exposure of Primary Human Muscle Cells to Plasma Fatty Acids Associated with Obese Phenotype Induces Persistent Suppression of Muscle Mitochondrial ATP Synthase beta Subunit. PLoS One 11, e0160057.

Tran L, Kras KA, Hoffman N, Ravichandran J, Dickinson JM, D’Lugos A, Carroll CC, Patel SH, Mandarino LJ, Roust L & Katsanos CS. (2018). Lower Fasted-State but Greater Increase in Muscle Protein Synthesis in Response to Elevated Plasma Amino Acids in Obesity. Obesity (Silver Spring) 26, 1179–1187.

Tran L, Langlais PR, Hoffman N, Roust L & Katsanos CS. (2019). Mitochondrial ATP synthase β-subunit production rate and ATP synthase specific activity are reduced in skeletal muscle of humans with obesity. Exp Physiol 104, 126–135.

Vanderboom P, Zhang X, Hart CR, Kunz HE, Gries KJ, Heppelmann CJ, Liu Y, Dasari S & Lanza IR. (2022). Impact of obesity on the molecular response to a single bout of exercise in a preliminary human cohort. 1091–1104.

Venojärvi M, Korkmaz A, Aunola S, Hällsten K, Virtanen K, Marniemi J, Halonen JP, Hänninen O, Nuutila P & Atalay M. (2014). Decreased Thioredoxin-1 and Increased HSP90 Expression in Skeletal Muscle in Subjects with Type 2 Diabetes or Impaired Glucose Tolerance. BioMed Research International 2014, 1–6.

Wada H, Kito K, Caskey LS, Yeh ET & Kamitani T. (1998). Cleavage of the C-terminus of NEDD8 by UCH-L3. Biochem Biophys Res Commun 251, 688–692.

Wisniewski JR, Zougman A, Nagaraj N, Mann M & Wi JR. (2009). Universal sample preparation method for proteome analysis. Nature Methods 6, 377–362.

Xia Q, Casas-Martinez JC, Zarzuela E, Munoz J, Miranda-Vizuete A, Goljanek-Whysall K & McDonagh B. (2023). Peroxiredoxin 2 is required for the redox mediated adaptation to exercise. Redox Biol 60, 102631.

Xu N, Gulick J, Osinska H, Yu Y, McLendon PM, Shay-Winkler K, Robbins J & Yutzey KE. (2020). Ube2v1 Positively Regulates Protein Aggregation by Modulating Ubiquitin Proteasome System Performance Partially Through K63 Ubiquitination. Circ Res 126, 907–922.

Yang S, Wang B, Humphries F, Hogan Andrew E, O’Shea D & Moynagh Paul N. (2014). The E3 Ubiquitin Ligase Pellino3 Protects against Obesity-Induced Inflammation and Insulin Resistance. Immunity 41, 973–987.

Yuan Y, Wang YY, Liu X, Luo B, Zhang L, Zheng F, Li XY, Guo LY, Wang L, Jiang M, Pan YM, Yan YW, Yang JY, Chen SY, Wang JN & Tang JM. (2019). KPC1 alleviates hypoxia/reoxygenation-induced apoptosis in rat cardiomyocyte cells though BAX degradation. J Cell Physiol 234, 22921–22934.

Zhao Y, Long MJC, Wang Y, Zhang S & Aye Y. (2018). Ube2V2 Is a Rosetta Stone Bridging Redox and Ubiquitin Codes, Coordinating DNA Damage Responses. ACS Cent Sci 4, 246–259.

